# Dynamic imaging of myelin pathology in physiologically preserved human brain tissue using third harmonic generation microscopy

**DOI:** 10.1101/2024.09.06.611764

**Authors:** Niels R.C. Meijns, Max Blokker, Sander Idema, Bert A. ’t Hart, Mitko Veta, Loes Ettema, Juliet van Iersel, Zhiqing Zhang, Geert J. Schenk, Marie Louise Groot, Antonio Luchicchi

## Abstract

Myelin pathology is known to play a central role in disorders such as multiple sclerosis (MS) among others. Despite this, the pathological mechanisms underlying these conditions are often difficult to unravel. Conventional techniques like immunohistochemistry or dye-based approaches, do not provide a temporal characterization of the pathophysiological aberrations responsible for myelin changes in human specimens. Here, to circumvent this curb, we present a label-free, live-cell imaging approach of myelin using recent advancements in nonlinear harmonic generation microscopy applied to physiologically viable human brain tissue from post-mortem donors. Gray and white matter brain tissue from epilepsy surgery and post-mortem donors was excised. To sustain viability of the specimens for several hours, they were subjected to either acute or organotypic slice culture protocols in artificial cerebral spinal fluid. Imaging was performed using a femtosecond pulsed 1060 nm laser to generate second harmonic generation (SHG) and third harmonic generation (THG) signals directly from myelin and axon-like structures without the need to add any labels. Experiments on acute human brain slices and post-mortem human slice cultures reveal that myelin, along with lipid bodies, are the prime sources of THG signal. We show that tissue viability is maintained over extended periods during THG microscopy, and that prolonged THG imaging is able to detect experimentally induced subtle alterations in myelin morphology. Finally, we provide practical evidence that live-cell imaging of myelin with THG microscopy is a sensitive tool to investigate subtle changes in white matter of neurological donors. Overall, our findings support that nonlinear live-cell imaging is a suitable setup for researching myelin morphology in neurological conditions like MS.

## Introduction

Myelin is classically seen as a lipid-rich passive electrical insulator, that prevents the leakage of current and allows for the efficient saltatory propagation of action potentials along axons. However, recent research has revealed that myelin may play a more complex role in neural function than previously considered, namely by the trophic support of energy-rich monocarboxylates (lactate and pyruvate) to energy-demanding axonal mitochondria [1]. The latter notion is particularly relevant to comprehend the pathological features of disorders like multiple sclerosis (MS), leukodystrophy and, more recently, temporal lobe epilepsy (TLE), which represent a spectrum of diverse conditions of unknown etiology, characterized by a fundamental implication of myelin aberrations [2].

A common feature of these disorders is the progressive disintegration and loss of myelinated axons in both the white (WM) and the gray matter (GM) in the brain, with lesions localized according to the specific pathological condition. Whether these axonal and myelin deteriorations are a mere consequence of other processes, or whether they represent a stand-alone etiopathological correlate of these diseases is still unknown. Recent evidence from our lab is in support of the latter possibility for MS, where the so-called normal appearing WM (NAWM) — regions of the MS brain not exhibiting signs of demyelination or overt inflammatory activity — presents with subtle myelin pathology. These morphological changes are characterized by a precarious detachment of myelin sheaths from their enveloping axonal and widening of myelin lamellae. This leads to the formation of typical myelin bulges (myelin blisters) detectable with immunohistochemistry in both post-mortem MS WM tissue and in the corpus callosum of relevant animal models of the disease [3–5]. Although the nature of this seemingly MS-specific pathological feature is still under investigation, a likely cause is the subtle instability at the level of the recently described axo-myelinic synapse (AMS), a glutamate-mediated form of communication between axon and myelin thought to support myelination and axonal action potential propagation [6,7]

To elucidate the factors contributing to these morphological changes in neurodegenerative conditions, it is essential to explore the cascade of pathophysiological and biochemical dynamics at their origin. Traditional methods like immunohistochemistry, despite their high-quality identification of such aberrations, do not offer insights into the dynamics of the pathophysiological process(es). On the other hand, the mere introduction of dyes into cellular systems for live-cell imaging may evoke unwanted alterations of biological processes [8]. Recent advancements in label-free live-cell imaging prove invaluable for examining these aspects in greater depth. Studies from our labs have already described the exploitation of a nonlinear microscopy approach by virtue of a second- and third harmonic generation (SHG, THG) microscope as a useful strategy to visualize with high-fidelity and high subcellular resolution, lipid-rich structures like myelin sheaths in both mouse and human specimens [9,10]. Moreover, we have shown that this data can be quantified and statistically evaluated [11,12] for clinical applications. Despite this progress, exploration into the crucial (patho)physiological changes in brain material connected to specific diseases has been hampered by the lack of a human brain model where cerebral tissue is kept viable for extended periods of time.

By combining an optimized protocol to preserve freshly excised post-operative and post-mortem human brain samples [13] with a custom-designed higher harmonic generation microscope, we explored the potential to visualize myelin alterations in acute human brain slices over time specifically using nonlinear microscopy. Furthermore, we extended our investigations employing a new organotypic slice culture model of human brain tissue which was recently published [14], to assess the longitudinal imaging capabilities of our setup.

In this study, we assessed the resolving power of the microscope setup, employed post-acquisition techniques to reconstruct complex myelin architecture and extended our setup to perform quantification of myelin vacuolization during unphysiological ion exposure, being a condition of the MS brain [15], and pharmacological inhibition of critical processes thought to be involved in MS axon-myelinic instability. In particular, in line with previous studies from our lab [5,16] and evidence on living MS subjects [15], here we tested whether a progressive increase of extracellular sodium concentrations evokes myelin swelling/blisters while imaging over time, and if the blockade of the calpain-cathepsin axis, shown to be involved in axon-myelin tethering protein degradation [17], may prevent such process.

Overall, our findings advocate that THG live-cell imaging represents a potential ideal 4D tracking system to investigate specific mechanisms underlying the formation of myelin pathology in human brain tissues, highlighting its significance for future research in other brain pathologies.

## Materials and Methods

### Laser-induced photodamage assessment

To assess photodamage induced by laser exposure to the tissue cells, human D384 medulloblastoma cells were cultured to 90-100% confluence in poly-L-lysine-coated dishes equipped with a 500 μm grid (μ-Dish 35 mm high, Grid-500, Glass Bottom, Ibidi) and incubated at 37°C. Following cultivation, cells were subjected to femtosecond-pulsed infrared (IR) laser irradiation under multiple settings. Post-irradiation, the cells were further incubated for 24 hours, then fixed with paraformaldehyde (PFA) and co-stained with Hoechst and propidium iodide (PI) at a concentration of 5µg/mL. This approach allowed for the delayed assessment of cell damage and the identification of irradiation-induced damage, which may not be immediately evident. The experiment was conducted in quadruplicate for every combination of laser pulse frequency and exposure duration. Control groups included cells unexposed to irradiation (negative control) and cells irradiated with UV light for 30 minutes (positive control). Specific cells within the grid were designated for each measurement condition to facilitate navigation. To prevent overlapping effects of irradiation, each targeted cell was isolated from neighbors that were not irradiated. In one set of experiments, pulse frequency ranged from 0.75 to 2.25 pulses per µs, with exposure times varying from 1.8 to 90 seconds. For exposure times exceeding 1.8 seconds, a 5-second intermission was introduced between exposures. The energy per pulse was maintained at a constant with the acousto-optic modulator (AOM) amplitude set to 2500, resulting in sample plane power levels between 2 to 12 mW. Experiments on photodamage were conducted in quadruplicate.

### Human Brain Samples

Human brain samples were obtained from 6 patients undergoing surgical epilepsy treatment at the Amsterdam University Medical Center (Amsterdam UMC). All patients provided written informed consent for tissue biopsy collection and signed a declaration allowing the use of their biopsy specimens in scientific research. Structurally normal brain tissue pieces, sizing approximately 1 × 1 × 0.5 cm, were excised from the temporal cortex and subcortical WM. The post-mortem data in this report are based on material received from individuals with MS and non-neurological disorders (non-MS). Samples with a post-mortem delay of less than 7 hours obtained from the Dutch Brain Bank were eligible for inclusion. For experiments using acute slices from post-mortem donors, immediately after resection, tissue samples were cooled for ten minutes in ice-cold N-methyl-D-glucamine (NMDG) buffer comprising: 93 mM NMDG, 2.5 mM KCl, 1.2 mM NaH2PO4, 20 mM HEPES, 12 mM N-acetyl-L-cysteine (NAC), 5 mM sodium ascorbate, 3 mM sodium pyruvate, 10 mM MgSO4, 30 mM NaHCO3, and 25 mM glucose; pH adjusted to 7.4 adapting a protocol previously published [18]. Using a surgical scalpel, slices were cut to create flat tissue surfaces. Protective slice recovery then commenced in oxygenated NMDG at room temperature for three minutes with carbogen (95% O2/5% CO2, at a 0.1 L/min flow rate), followed by a 60-minute oxygenated incubation at room temperature in 50 mL of HEPES holding solution, containing: 92 mM NaCl, 2.5 mM KCl, 1.2 mM NaH2PO4, 20 mM HEPES, 1 mM NAC, 5 mM sodium ascorbate, 3 mM sodium pyruvate, 0.5 mM MgSO4, 1 mM CaCl2, 30 mM NaHCO3, and 25 mM glucose; pH adjusted to 7.4. Subsequently, each slice was placed on a #1.5H glass coverslip (µ-Dish 35 mm, high glass bottom, ibidi, Gräfelfing, Germany) for live-cell recording, secured with a custom-cut sponge or harp to minimize tissue drift during imaging. For all experiments with human tissue, we adhered to the Netherlands Code of Conduct for Research Integrity and the Declaration of Helsinki, with protocol approval by the Medical Ethical Committee of the Amsterdam UMC. Human samples were collected from first of January 2021 to 2024.

### Cortical slice cultures

For experiments employing cultured human brain tissue, after resection, a segment of the middle frontal gyrus was collected in ice-cold Hibernate-A (Invitrogen, A12475-1, Waltham, Massachusetts) supplemented with 1% Penicillin-streptomycin (Invitrogen, A15140-122). The meninges and blood vessels were removed, and a ∼1.5 x 1.5 cm block containing equal parts of cortex and white matter was affixed with superglue and supported by agar gel on a support platform. Following submersion in dissection medium, 400 µm slices were cut with a vibratome and transferred to a membrane culture insert (Sigma-Aldrich, PICMORG50, St. Gallen, Switzerland). The culture media consisted of 50% MEM (Invitrogen, 32360-026), 25% Earle’s Balanced Salt Solution (Invitrogen, 24010-043), 25% Heat-inactivated Horse Serum (Invitrogen, 26050-088), 1% GlutaMAX-I (Invitrogen, 35050-038), 1.4% of 45% D-Glucose (Sigma, G8769), 1% Penicillin-streptomycin (Invitrogen, 15140-122), and 0.5% Amphotericin B (Invitrogen, 15290-018). The media was filter sterilized using a 0.22 µm filtration unit. Culture media was changed 24 hours after slicing and subsequently every two days. The protocol was adapted from previously published and collaborated work from Bugianís group [19].

### Third harmonic generation microscopy

Higher harmonic generation is a nonlinear scattering process resulting from femtosecond laser light interaction with tissue molecules. Signal generation may occur in tissue depending on phase matching conditions and nonlinear susceptibility coefficients. The third order harmonic process results from the combination of three photons from the incident laser beam into one photon with triple the energy and thus one-third of the excitation wavelength, via a virtual energy state. Utilizing a laser wavelength in the near-infrared spectrum results in generation of the harmonic signal in the visible light range. The intensity of THG production is given by the equation:

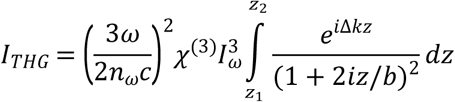

with laser intensity 𝐼_𝜔_, angular frequency 𝜔, medium refractive index 𝑛_𝜔_, speed of light 𝑐, phase mismatch Δ𝑘 = (𝑛_3𝜔_3𝜔/𝑐) ― 3(𝑛_𝜔_𝜔/𝑐), position along the beam axis 𝑧, boundaries of the medium 𝑧_1_ and 𝑧_2_, and the confocal parameter of the focused laser beam 𝑏. Optimal THG production occurs when a small inhomogeneity is introduced in the laser focus, or at structural interfaces.

Imaging of the brain tissue slices was performed on an inverted THG microscopy setup. Based on the susceptibility of the material, 𝜒^(3)^ and phase-matching conditions Δ𝑘, a THG signal can be produced. Brain tissue is a suitable candidate for THG imaging due to the lipid-rich components acting as the major contrast source. In typical images of structurally normal white matter, most brain cells appear as dark holes while myelinated axons are clearly visible. The system employs a femtosecond laser (SL-F300, Seed Lasers, Lithuania) emitting 100 fs pulses at 1060 nm, with a maximum output of 20 nJ per pulse. Generated signals are collected in a backscatter configuration using an oil-immersion objective (40×/1.3 NA, Nikon). Backscattered THG photons at 1/3^rd^ of the laser wavelength are split from the fundamental laser beam by a dichroic mirror (BrightLine FF872-Di01-25x36x2.0, Semrock, Inc.), filtered by a narrow-band interference filter (BrightLine F01-355/40-25, Semrock, Inc.) and collected with a high-sensitivity photomultiplier tube (H10721-210, Hamamatsu Photonics K.K.). An acousto-optic modulator (MT250-A0.5-1064, AA Opto Electronic), controlled by a radio frequency driver (MODA250, AA Opto Electronic), is situated directly after the laser shutter to modulate the duty cycle and amplitude.

### THG imaging of myelin

Sample holders with prepared brain slices were transferred to a stage-top incubation chamber (H301-PRIOR-H117, OKOLAB S.R.L.) and maintained at a constant temperature of 37 °C under continuous carbogen (95% O2/5% CO2) flow. For live-cell imaging, continuous perfusion with artificial cerebrospinal fluid (aCSF) was conducted at a rate of 1 mL/min using peristaltic pumps (ISM832C, ISMATEC, Cole-Parmer GmbH). The aCSF composition is as follows: 125 mM NaCl, 15 mM KCl, 2 mM CaCl, 1.25 mM NaH_2_PO_4_, 1 mM MgSO_4_, 26 mM NaHCO_3_, 10 mM C_6_H_12_O_6_; with a pH maintained at 7.4. Additionally, the setup was designed to allow for sample perfusing without inducing tissue drift or compromising image quality. For our experiments, we utilized both forward and backward detection mechanisms of higher harmonic generation signals, to capture comprehensive images of myelin sheaths. Notably, backward-detected THG signals were exclusively analyzed in our photodamage studies. This approach facilitates 4D data acquisition through time-lapse imaging of 3D depth scans. For identifying a suitable region-of-interest (ROI), the scanning software (Flash Pathology BV) was used in inspection mode to scan the tissue across a 400 × 400 µm field-of-view (FOV) at an acquisition speed of 2 frames per second, employing bi-directional scanning within a 400 × 400-pixel image. The motorized stage (Prior Scientific) enables control over the x, y, and z movement with a minimum step size of 0.2 µm. Time-lapse recordings were conducted in scanning mode, using either a 200 × 200 µm or 400 × 400 µm FOV within a 1000 × 1000-pixel image at an acquisition speed of one frame per 1.8 seconds. To mitigate water heating due to light absorption, a cooling-down period of 5 minutes was introduced between recordings.

Recordings, including THG time-lapse sequences, were initially saved as 8-bit grayscale BMP files and then automatically processed for histogram normalization before being converted into TIFF stacks using the Fiji software suite for further data analysis [20].

### THG experiments on corpus callosum tissue

Acute slices excised from genu/corpus of the corpus callosum of both MS and non-MS control donors were treated as mentioned above and exposed to increasing concentrations of aCSF [Na^+^], from 125 mM to 133 and later 140 mM) after acquisition of a minimum of 3 baseline z-stacks during 125 mM [Na^+^]. To induce an opening of sodium voltage channels in axons, the concentration of KCl is aCSF was increased from 2.5 mM to 15 mM for at least 1 hour. Donor material was exposed to increasing concentrations of NaCl in presence ALLN (20 uM, Sigma-Aldrich, Germany) to test whether the blockade of this pathological process was able to inhibit swelling formation in MS WM.

**Table 1.** Demographic and clinical Information of human MS and healthy control (HC) samples.

| Case | Sex | Age (years) | pH CSF | Diagnosis | PMD (HH:MM) | COD |
| --- | --- | --- | --- | --- | --- | --- |
| C17 | F | 75 | N/a | MS | >7 | N/a |
| C18 | M | 58 | 6.07 | MS | 08:20 | Euthanasia |
| C20 | N/a | N/a | N/a | HC | >7 | N/a |
| C21 | N/a | N/a | N/a | HC | >7 | N/a |
| C28 | F | 44 | 6.64 | MS | 05:55 | End-stage MS |
| C30 | F | 80 | N/a | MS | 05:35 | Euthanasia |
| C35 | F | 90 | 6.6 | HC | 03:40 | Euthanasia |
| C36 | M | 78 | 6.2 | MS | 06:55 | Dehydration |
| C37 | M | 92 | 6.1 | HC | 07:45 | Lung fibrosis |
| C38 | M | 86 | N/a | HC | >7 | N/a |
| C39 | M | 91 | 6.83 | HC | 06:35 | Heart failure |

CSF: cerebrospinal fluid, PMD: post-mortem delay, COD: cause of death.

### Identification and quantification of myelin swellings

To semi-automatically quantify the density of swellings in THG z-stack time lapses we first registered and stabilized the images to counter any drift observed during imaging. Z-stacks were subsequently inverted, to highlight non-myelin, and median filtered (2 px). Histograms of THG z-stack were equalized with contrast-limited adaptive histogram equalization (CLAHE). A probability mapping for non-myelin signal was then created by sparse annotation in Ilastik. Possible swelling objects were then detected from non-myelin probability maps by thresholding (core: 0.85, final: 0.5) and size filtering. Sufficiently round objects were marked as myelin swellings using the Ilastik object detection module [21].

### Histological myelin staining

To correlate myelin observed with THG imaging with conventional histology a myelin staining was conducted on human brain sections using a 0.1% solution of Luxol Fast Blue (LFB; Gurr, UK), incubated in an oven at 58°C overnight. Subsequent washing steps involved brief immersion in 96% ethanol for 2-3 seconds, followed by rinsing in Milli-Q (MQ) water for 3 seconds. Differentiation was then achieved by treating the sections with a 0.05% lithium carbonate solution (Merck Millipore, Germany) for 5 seconds, and subsequently, with 70% ethanol for 5-7 seconds, until the gray matter became colorless and the white matter retained the blue hue. The sections were rinsed again in MQ water, dehydrated through a graduated alcohol series (96% for 3-5 minutes, then 100% for two 5-minute periods), cleared in xylene (for three 5-minute periods), and finally mounted with Entellan under a coverslip. Normal-appearing white matter (NAWM) was characterized as regions exhibiting typical levels of proteolipid protein (PLP) immunoreactivity distributed evenly across white matter areas, with an absence of major histocompatibility complex class II (MHC-II) immunoreactivity [22].

### SHG imaging of axon-like structures

To test the ability of our setup in visualizing axonal processes abreast the THG visualization of myelin human neuroblastoma SH-SY5Y cell cultures, following differentiation, were prepared on poly-L-lysine-coated dishes featuring a grid (μ-Dish 35mm, high, Grid-50, Glass Bottom, from Ibidi). These cells underwent a 7-day cultivation process at 37°C, with the culture medium being replaced several times. This medium included retinoic acid (RA) for the initial four days to initiate differentiation, followed by human brain-derived neurotrophic factor (hBDNF) for the remaining three days to further promote neuronal differentiation. Subsequent to the 7-day incubation and differentiation period, the cultures were subjected to imaging using a Second Harmonic Generation (SHG) imaging setup. A bidirectional scanning protocol was employed with a laser exposure time of 1.8 seconds and a field of view (FoV) of 200 x 200 μm. On average, five acquisitions were made for each measurement. The setup was adjusted to emit 2.25 pulses per microsecond (μs) for the majority of measurements. The output power at the level of the sample varied across measurements but maintained an average of approximately 4 mW.

### Immunohistochemistry

Cells and tissue were fixed in 4% paraformaldehyde (PFA) for 30 minutes. After fixation phosphate-buffered saline (PBS, Sigma D8537) was used to rinse the cells and cultures, which were then stored in the refrigerator until staining. Staining for microtubule-associated protein 2 (MAP2) and neurofilament-light (SMI312) utilized primary antibodies diluted at 1:1000 in 2% normal serum in PBS, after initial blocking and permeabilization steps with normal serum and Triton X-100, respectively. This was followed by incubation with Alexa Fluor conjugated secondary antibodies (488 for MAP2 and 546 for SMI312) at a 1:400 dilution, for 1h in the dark. Fluorescence from the Alexa Fluor conjugated antibodies was observed under a Nikon ECLIPSE Ti2 inverted microscope, employing a 40x oil immersion objective (1.3 NA) across a 442.20 x 442.20 μm FoV. Precise location mapping, utilizing the dish grid, enabled correlation of sites imaged with the SHG setup. The MAP2 signal was detected with a FITC filter block (ex. 470 nm, em. 530 ± 21.5 nm), while SMI312 utilized a Texas Red filter block (ex. 555 nm, em. 630 ± 34.5 nm).

## Results

### THG microscopy does not induce nonlinear photodamage in human D384 medulloblastoma cells

Photodamage during nonlinear imaging, particularly in live cell studies, remains a significant concern as it can lead to artifacts and convolution of biological processes [23,24]. Long-term imaging capacity is essential for dynamic and developmental studies [25]. Within this context, THG microscopy inherently conserves energy, as no photons are absorbed or emitted. Nonetheless, its requirement for high irradiation and the consequent involvement in absorption-based phenomena such as two-photon fluorescence (2PF), alongside potential issues like the self-absorption of emitted THG light and the generation of phototoxic substances, raises concerns about potential photodamage. To this end, we aimed to systematically evaluate the extent of photodamage associated with THG imaging sessions. Human D384 medulloblastoma cells grown in confluent cultures were irradiated with sequences— ranging in pulse frequencies from 0.75 to 5.25 pulses per µs and exposure times from 1.8 to 90 seconds. Quantification of cell damage indicated no clear non-linear cell damage in the tested range of pulses per microsecond and exposure time (Fig 1A). Longer exposure times do indicate a slight trend toward more photodamage in the 5.25 pulses/µs condition, although no statistically significant differences were observed between 1.8 s and 90 s irradiation time for the 5.25 pulses/µs condition. Induced photodamage with 90s continuous irradiation at 5.25 pulses/µs was most pronounced at the center (Fig 1). We conducted subsequent experiments using a constant energy per pulse (∼4.5 nJ per pulse) combined with 3.75 pulses/µs for sufficient THG signal.

**Fig 1:**
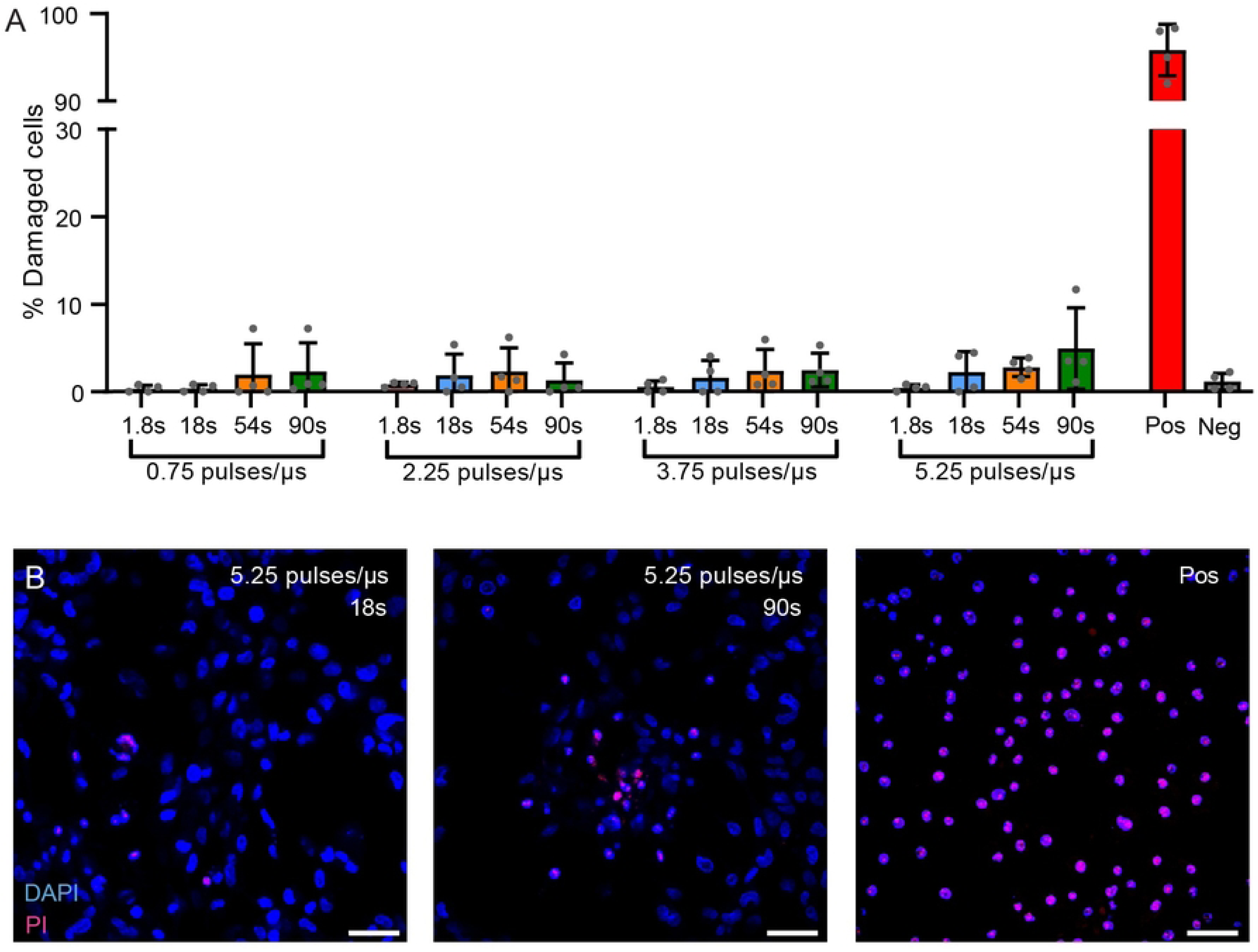
Photodamage is in human D384 medulloblastoma after THG imaging. (A) The percentage of photodamaged cells following various laser pulse frequencies (0.75 to 5.25 pulses/µs) across exposure durations (1.8 s to 90 s), with controls for baseline (Neg) and UV-induced damage (Pos). No effects were observed for pulses/µs with a one-way ANOVA (F(3,12) = 0.982, p = 0.44). In the most extreme condition (5.25 pulses/µs) no significant difference in in cell damage was observed between 1.8s and 90s (t(6) = 1.921, p = 0.1031). (B) Representative fluorescence images corresponding to the highest pulse exposure condition (5.25 pulses/µs) at 18 s and 90 s, with DAPI (blue) marking nuclei and PI (red) indicating damaged cells. For all conditions consisting of more than one accumulation (an exposure time longer than 1.8 s), a 5-second interval was applied between subsequent exposures. The energy per pulse was kept constant and was estimated to be ∼4.5 nJ per pulse based on power measurements at the sample plane. Error bars indicate SD. Scalebars are 50 µm. α = 0.05.

### Visualization of myelinated axons in physiologically viable human brain slices using THG

After determining the safe margins for THG imaging without inducing photodamage while maintaining signal quality, we set out to image myelin in human brain tissue. Human brain slice cultures were isolated and kept in culture until DIV12 and prepared for imaging. THG imaging of these cultures enables production of a three-dimensional cutaway from a z-stack image, revealing intricacies within the cortical gray matter with clear detail (Fig 2A). Striking similarities were observed between THG and Luxol Fast Blue (LFB), a common histological dye for myelin (Fig 2B). Both THG and LFB capture subtle myelin pathologies, like myelinic swellings. Significant variations in THG signal-to-noise (S/N) ratios were noted in deeper tissue layers during THG imaging, primarily attributed to signal scattering and optical aberrations within the sample, culminating in focal misalignments of the laser excitation. These discrepancies were notably more prominent in WM tracts compared to GM regions, consequently affording superior imaging quality at increased tissue depths within the gray matter (Fig 2C). Intriguingly, despite the ubiquity of lipid bilayers within human brain architecture, not all lipid interfaces produced detectable THG signals. In the assessment of the microscopy system’s optical performance, the spatial resolution was a critical parameter. To quantify the radial resolution, we utilized a broken glass capillary as a sample. Fig 2D illustrates the approach through an intensity plot and edge-spread function derived from a line measurement across the THG image of the glass capillary. We estimated the Full Width at Half Maximum (FWHM) to be approximately 400 nm for lateral resolution (Fig 2D). Additionally, the axial resolution, assessed similarly, through imaging a glass bottom dish, was found to be roughly 1300 nm (Fig 2E).

**Fig 2:**
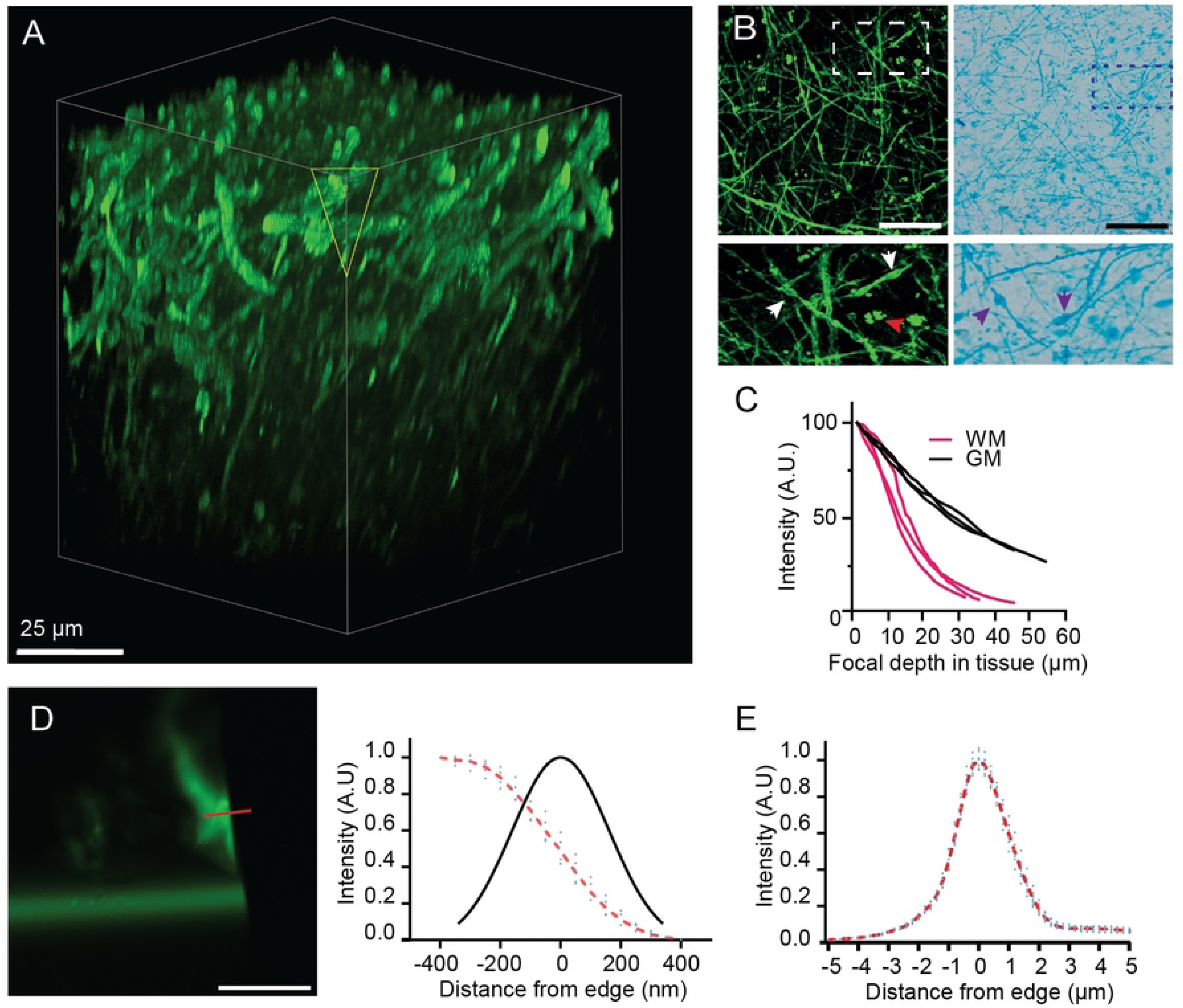
Optical Performance of nonlinear microscopy setup. (A) A three-dimensional cutaway from a z-stacked THG image. Volumetric rendering (100 × 100 × 100 µm) (THG) image highlighting cortical GM, showcasing the technique’s capability to provide detailed structural insights. (B) Comparative analysis of myelin imaging using THG against Luxol Fast Blue (LFB) staining of cortical tissue. Insets highlight myelinic swellings visible in both THG and LFB images, with THG’s sensitivity to lipid bodies (red arrow in THG inset) and swelling (white and blue arrowheads). (C) Comparison of the relation between THG signal intensity relative to the imaging depth from the tissue surface, comparing gray and white matter regions (n=3), illustrating THG signal properties differences between the regions. (D) THG image captured at the edge of a glass capillary, accompanied by a superimposed line-scan analysis (left). Intensity profile across a redline and the corresponding line spread function, showcasing the system’s spatial resolution as measured from the capillary edge (right). (E) Full width at half maximum (FWHM) for axial resolution is approximately 1.3 μm, determined through imaging a glass bottom well. Scale bars is 15 μm, unless specified otherwise.

### Subtle myelin pathology can be monitored with time-lapse THG microscopy

To test whether our tool captures subtle myelinic changes over-time in human tissue, a human cortical slice culture (DIV12) was prepared for imaging in a 35 °C custom imaging chamber perfused with carbogenated artificial cerebrospinal fluid, extending a previously published protocol [19]. To demonstrate the longitudinal monitoring capabilities of our experimental setup in tracking myelin pathology using THG microscopy while maintaining stringent experimental controls, we conducted time-lapsed z-stacks (Fig 3A). Notably, subcortical U-fibres, which interconnect adjacent regions within cortical gyri, exhibited finely ensheathed myelin structures after twelve days in vitro. Upon exposure to 1 mM sodium azide and 0.2 mM glutamate, distinctive membrane irregularities in the myelin sheaths manifested within a five-minute timeframe (Fig 3B). Changes in THG intensity at the membrane interfaces contrasting their immediate microenvironment were distinctly visible along the axonal length. Furthermore, myelin swellings or blister formations were observed at various axonal loci, with these swellings progressively increasing in size throughout the observation period. By analyzing cross-sectional views of axons alongside corresponding profile plots, we could effectively discern abaxonal and adaxonal myelin membranes, allowing for g-ratio calculations (Fig 3C).

**Fig 3:**
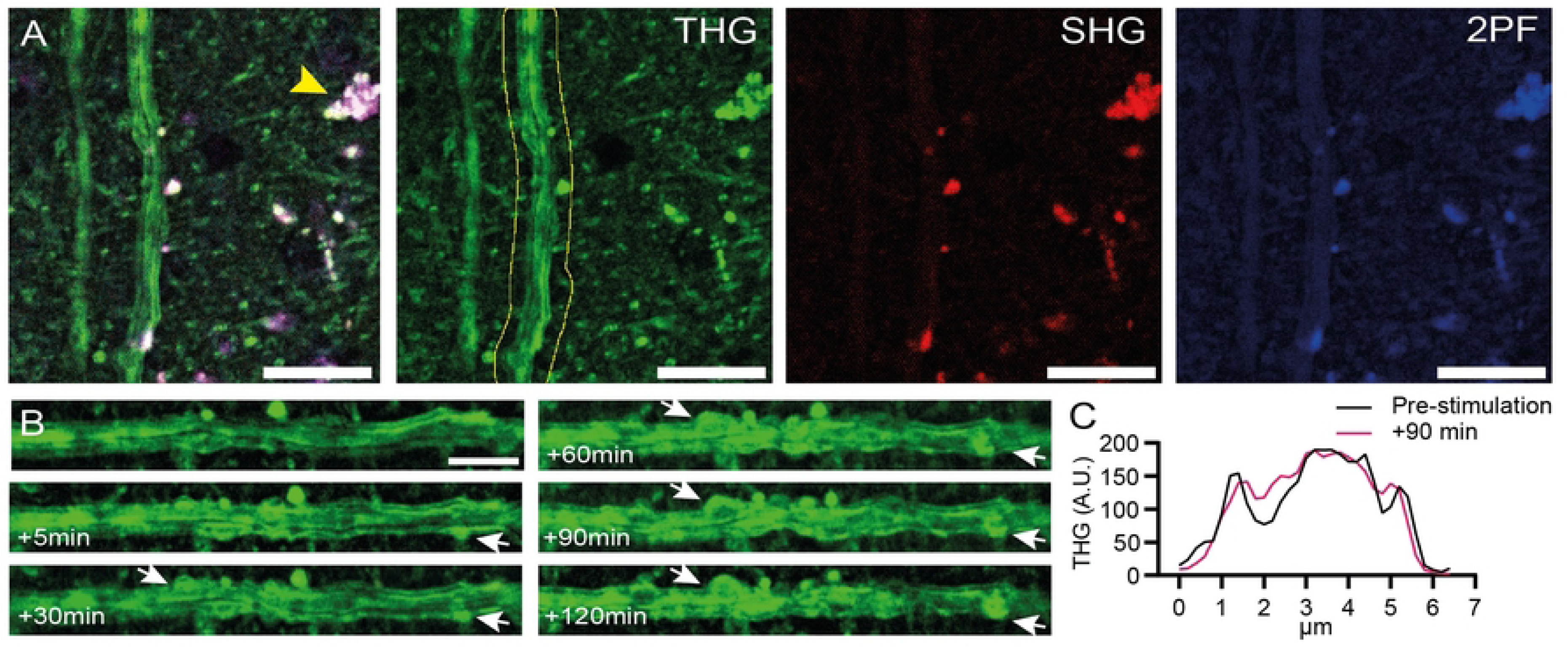
Subtle alterations in myelin morphology during THG imaging. (A) Example of a myelinated axons in human cortical slice culture (DIV12). Lipofuscin in post-mortem neuronal soma is broadly fluorescent across SHG (red at 525nm) and 2-photon (blue at >580 nm) channels (arrowhead). (B) Swellings in myelin sheath develop post 1 mM sodium azide and 0.2 mM glutamate stimulation, persisting during 2-hour acquisitions (white arrows indicate). (C) Post-stimulation, myelin membrane demarcation (ab- and adaxonal) becomes indistinct as shown in the profile plot (line not depicted). Scalebar A is 20 µm. Scalebar B is 10 µm.

### Quantification and inhibition of myelin swelling formation

To further showcase the suitability of our setup for longitudinally imaging of myelin we set out to probe human corpus callosum of MS patients and non-MS patients during time lapsed THG imaging. Myelin pathology was induced by administration of increasing sodium ([Na+]) concentrations aiming at overstimulation of the axon myelinic synapse and to induce myelinic blisters/swelling [3]. To that end, we increased the [Na+] in the perfused aCSF during imaging in stepwise fashion with 7mM increments after approximately 30 and 120 minutes respectively (Fig 4A). We utilized a machine-learning-based approach to volumetrically segment myelin and subsequently classify myelic swellings in z-stack THG acquisitions (Fig 4C). Interestingly, post-mortem resected corpus callosum tissue was considerably more riddled with myelin swellings compared to non-MS controls (3.17 x 10^5^ ± 1.6 x 10^5^ per mm^3^ MS vs 0.64 x 10^5^ ± 0.17 x 10^5^ non-MS). The difference was found to be statistically significant (t(5)= 3.068, p = 0.02). Increasing [Na+] did not show statistically significant difference between MS and non-MS WM (F(1.196, 7.173) = 2.978, p = 0.124). To test whether myelinic swelling formation could be prevented by limiting protease activity, we preincubation tissue sections with N-Acetyl-L-leucyl-L-leucyl-L-methioninal (ALLN), a peptide known to inhibit calpain, cathepsin L and cathepsin B. Preincubation did result in annulment of the swelling increase seen in MS corpus callosum (Fig 4D) for all timepoints (t(18)= 3.141, p = 0.005).

**Fig 4:**
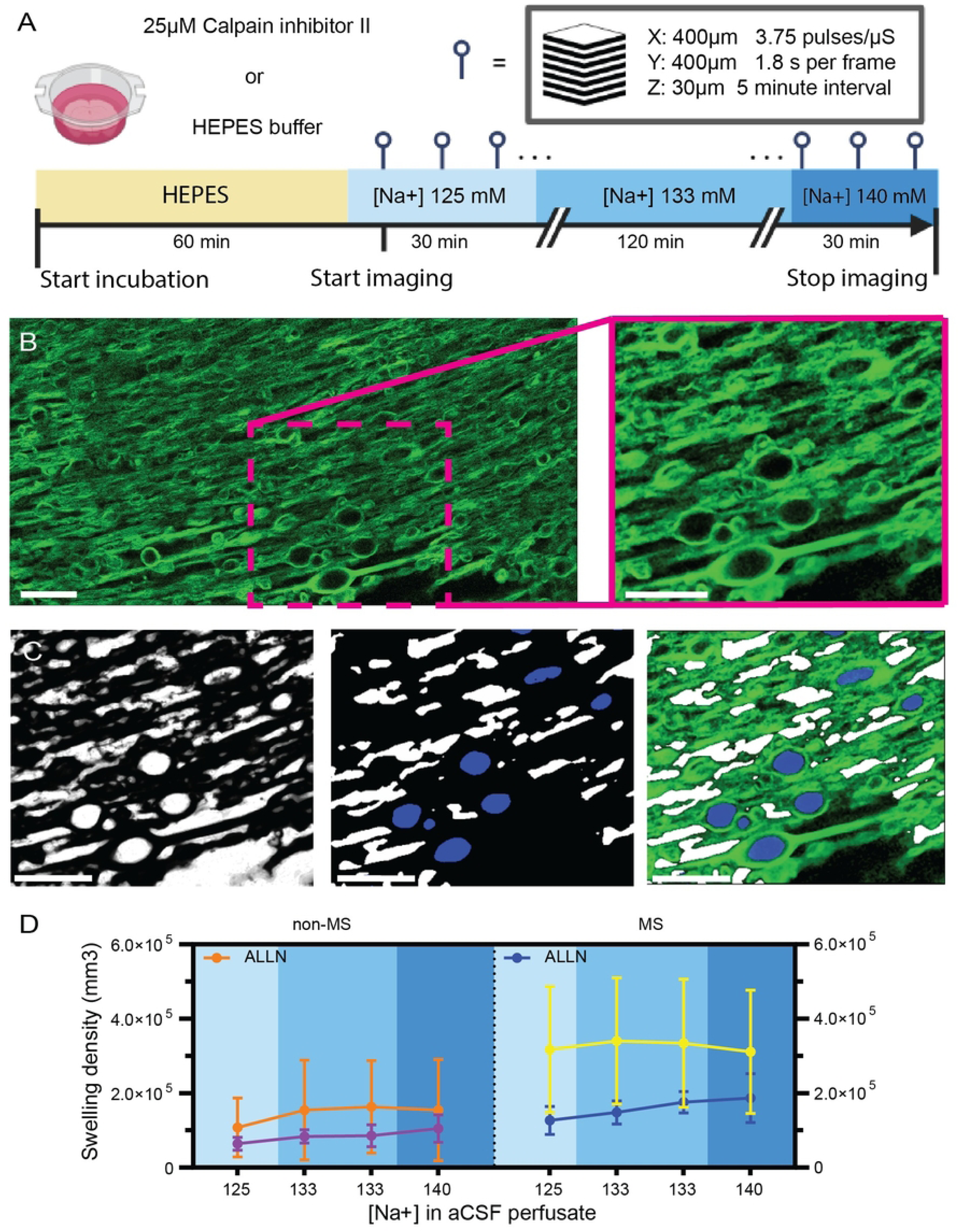
Inhibition of calpain-cathepsin axis with ALLN annuls the development of myelin swellings in MS corpus callosum. (A) Schematic illustrating the treatment of corpus callosum slices with N-Acetyl-L-leucyl-L-leucyl-L-methioninal in HEPES buffer followed by exposure to artificial cerebrospinal fluid (aCSF) adjusted to various [Na+] concentrations to simulate conditions that may induce myelin swelling. (B) Representative THG micrograph of myelin during imaging session. Myelinic swellings can readily be observed. Inset: THG signal after CLAHE contrast enhancement. (C) Inverse pixel-level probability map of CLAHE-corrected THG signal (left). Middle: Object-level classification discerned myelinic swellings (blue) from extracellular space (white). Right: Composite visualization merge of the original THG signal imagery with the superimposed object identification data. (D) Quantitative analysis of swelling density across different sodium concentrations in aCSF for non-MS and MS corpus callosum. Scalebars in B are 20 µm. n=6 (non-MS) and n=5 (MS).

## Discussion

Methods to visualize myelin without altering the tissue constituents, adding dyes or destroying the tissue are limited. This represents a limitation for live-cell, high resolution imaging investigations critical to provide crucial information of the pathophysiological changes in a number of myelin-related conditions. Here, we demonstrate label-free multiphoton live-cell imaging of myelin in its white matter context by THG microscopy. THG has historically been used to image transparent tissues such as embryonic tissues, plant tissues, and *in vivo* imaging of small organisms [26]. Only more recently, this technique has found applications in instant pathology assessment of surgically resected cancerous tissues [9]. The main challenge in the preparation of acute human brain slices and organotypic slice cultures for THG imaging is recreating and maintaining the *in-vivo* physiological environment. To the best of our knowledge, no protocols for preparation of live human brain sections for THG microscopy exist thus far. As such, our report serves a platform to perform investigation on myelin morphology in live human brain tissue while allowing experimental control.

The time-to-autopsy interval is by far the most decisive factor affecting tissue viability and health. To minimize tissue decay upon retrieval, we used a combination of NMDG- and HEPES-based cutting and holding solutions aiming to avoid degenerative processes like Ca2+ excitotoxicity [27]. Although cutting tissue with a microtome may be less harmful to tissue integrity compared to other techniques such freehand cutting, cell damage at the cutting surface can obviously not be avoided. Imaging for (sub-)cellular morphological features required therefore a certain minimal tissue depth, to be able to create an image at location relatively unaffected by artefacts from tissue preparation. Using post-mortem human brain material offers numerous advantages compared to resected tissue; it allows for the collection of tissues from various brain regions, from matching controls, and from individuals with different neuropathologies.

Utilizing NIR wavelength photons to generate THG signal allows for image acquisition in deeper planes, although the main limiting factor in image quality was scatter of THG in the UV range. Previous attempts to quantify photodamage in non-linear optics have already determined the key factors for imaging without damage to be fast repetition rate, NIR excitation and short pulse duration [28]. Despite optimization of the excitation light, emitted UV-range THG light may be a concern, as it may produce reactive species. We observed no increase in cell damage upon overnight incubation under a wide range of settings able to produce THG, suggesting the effect of emitted UV light in the tissue itself is negligible. Interestingly, limited photodamage we observed was often in the center of the field of view, suggesting heat development as a likely culprit of cell damage [28]. As such, we confirm that high peak powers, while maintaining low average power with pulse repetition rate around 1-10 Mhz are optimal for long-term, life-cell, label-free imaging of myelin with THG microscopy.

The current THG imaging analysis does not distinguish between the previously documented swelling types [3]. Hence, we use the term ‘swelling’ throughout the paper to describe myelin abnormalities. We posit that discrimination between the various forms of swellings and their implications on myelin integrity may be essential for elucidating the pathophysiology of myelopathies. By characterizing these morphological changes at a subcellular level, we aimed to uncover novel insights into the mechanisms underlying myelin disintegration. Myelinated tracts of cortical u-fibres in cortical slice cultures could be imaged with THG for hours and quickly developed swellings upon exposure to glutamate and sodium azide. Since central myelin is known to be subject to axon-bourne glutamatergic injury following hypoxia, an event mediated by NMDA receptors, an interpretation of this finding might be ascribed to overstimulation of the axon-myelinic synapse [29,30]. In contrast, experiments on acute slices where the extracellular [Na^+^] was altered failed to report a swelling increase in MS tissue when compared with non-MS material. Possible explanations should consider factors such as i) the insufficient flux of Na^+^ into the axonal compartment after moderate increase of this ion in the aCSF (15 mM) or ii) the already enriched presence of swellings in baseline condition in MS material. The latter may be a direct effect of the enhanced prevalence of myelin blisters present in MS NAWM tissue [3].

Excessive Na+ inflow in axonal compartments was previously shown to instigate a reversal of Na+-Ca2+ exchanger with consequent increase of Ca2+ levels in MS lesional axons [31]. This effect was believed to overstimulate the AMS glutamate-release from axon to myelin and activate a cascade of pathological mechanisms that culminates in the activation of the calpain-cathepsin axis. The latter putatively elicits degradation of the myelin associated glycoprotein and promote myelin blistering and swelling [16]. One may argue that to induce such massive changes in AMS physiology a sustained alteration of Na+ homeostasis needs to take place. As a matter of fact, Na+ imaging on living MS volunteers showed higher total Na+ concentration in NAWM in comparison with healthy individuals [15]. Irrespective, preincubation of corpus callosum acute slices with ALLN, a compound designed to selectively block the Ca2+-dependent calpain-cathepsin axis, showed a substantial reduction of swellings in MS tissue from the very early time points (and throughout all the imaging time). Such effect was not observed in the non-MS samples. In conclusion, we demonstrate that label free imaging of myelin with THG has great optical properties (in 3D) and produces negligible photodamage. We imaged acute changes in myelin structure after chemical asphyxiation and demonstrate that MS brains present with more myelin swellings, which was annulled by preincubation with calpain inhibitor. Overall, given that myelin structure and organization is of interest in any kind of myelinopathy, our results provide a novel approach to investigate myelin through dynamic imaging.

## Acknowledgements

We thank Jeroen Geurts (Vrije Universiteit Amsterdam) for motivating support in the conceptualization of the project. We thank Martin Slaman, Tim Heistek (Vrije Universiteit Amsterdam) and Frank van Mourik (Flash Pathology B.V.) for assistance in the technical setup. We thank Irene Frigerio, John Bol, Allert Jonker, Marije Zwaneveld, Iris van den Heuvel, Annabel Timmers, Gema Muñoz González, Gaia Bonfiglio (Amsterdam UMC) for experimental assistance. We thank Fernando Nobrega Santos, Marko Popvic, Rebecca Vischjager, Chloé Adel and Loek van Steijn (Amsterdam UMC) for assistance in data analysis.

This study has been funded by InstantPathology project of the research program Applied and Engineering Sciences (nr: 15825, coordinated by MG), Nationale Wetenschapsagenda of the Dutch Research Council (NWO, Idea Generator nrNWA.1228.191.204 awarded to AL), Stichting MS Research, MoveS, Klimmen tegen MS (MS16-945b, awarded to JJG), National MS society (Progressive MS alliance award, PA2021-36033 awarded to AL) Dutch National MS Foundation (Research grant OZ2021-008 awarded to AL), Stichting Amodo, KNAW (awarded to JJG) and by Stiching Leef! (awarded to AL).

---

Fig S1: Detection of SHG signal in SH-SY5Y neurites.

SHG imaging of SH-SY5Y cells revealed, fibrillar structures, consistent with the development of ordered neural processes in their outgrowing neurites. The 2.25 pulses per microsecond exposure resulted in dim SHG signal during live imaging (Fig S1 A). Post-processing of accumulated frames subsequently enabled visualization of the neurite architecture without observable photodamage. Subsequent correlative immunohistochemistry validated the presence of mature axonal markers (Fig S1 B). Microtubule associated protein 2 (MAP2) and neurofilament-light immunofluorescent staining yielded clear overlap with transmitted SHG signal.

